# The Flash Mode of Proton Irradiation in the Bragg Peak Partly Spares the Embryogenesis of the Quail

**DOI:** 10.1101/2024.01.27.577528

**Authors:** Sergey V. Akulinichev, Segrey I. Glukhov, Yury K. Gavrilov, Dmitry A. Kokontsev, Elena A. Kuznetsova, Valeriia V. Martynova, Ivan A. Yakovlev

**Affiliations:** Institute for Nuclear Research, Russian Academy of Sciences, Troitsk (Moscow), 108840 RF; Petrovsky National Research Centre of Surgery, Moscow, 119991 RF; Institute of Theoretical and Experimental Biophysics, Russian Academy of Sciences, Pushchino, Moscow region, 142290 RF

**Keywords:** irradiation with protons, FLASH radiotherapy, quail embryos, irradiation of eggs

## Abstract

The study of biological effects of high-dose-rate ionizing radiation using animal models remains an urgent problem. In the work, a comparison of proton-induced damage to embryos (the 9^th^ post-fertilization day) and 1-day-old baby chicks (the 18^th^ post-fertilization day) from irradiated eggs of Japanese quail (*Coturnix coturnix japónica*) irradiated in and outside the Bragg peak with the conventional and ultra-high dose rate D (CONV D < 1 Gy/s, FLASH D ∼100 Gy/s and single-pulse flash - SPLASH D ∼10^6^ Gy/s, respectively) was carried out. The following criteria were used: survival, body weight and body length of embryos, weight of chicks, percentage of erythrocytes with anomalies (with micronuclei, without nuclei, with two nuclei), and the speed of movement of 1-day-old chicks in the open field test. The largest number of dead embryos was recorded after irradiation with 7 and 14 Gy in the plateau region of the proton trajectory. By the criteria of body weight and length, as well as the number of erythrocytes with micronuclei in 9-day-old embryos from eggs irradiated with a dose of 8.5 Gy in the Bragg peak, FLASH and SPLASH modes were found to be the least traumatic compared with the CONV mode. The weight of chicks from eggs irradiated with a dose of 4 Gy in the Bragg peak did not differ from the control and did not depend on the irradiation mode. The lowest death incidence and the smallest number of abnormal erythrocytes were recorded after FLASH and SPLASH irradiation; in chicks that hatched from eggs irradiated in the CONV mode, a tendency for an increase in the number of abnormal erythrocytes was observed. The speed of movement of chicks from FLASH- and SPLASH-irradiated eggs was comparable with that from unirradiated eggs, and chicks from eggs irradiated in the CONV mode were less active than chicks from unirradiated eggs and eggs irradiated in the other regimes. Thus, the irradiation of eggs with a beam of accelerated protons in the Bragg peak in the FLASH/SPLASH modes is less damaging for healthy tissues and for the development of embryos and chicks on the cellular, anatomical, and physiological levels.

## 1. Introduction

FLASH irradiation, as compared with the conventional irradiation (CONV, D < 1 Gy/s) is carried out with a high dose rate (D) > 40 Gy/s [1, 2]. Recent investigations confirmed broad potentialities of the FLASH technique, which uses electrons, photons, and protons to diminish the toxicity in healthy tissues in the brain, lungs, intestines, blood, bones, muscles, and skin of animals without reducing the tumoricidal activity of ionizing radiation [3]. In particular, some experiments demonstrated a better preservation of peripheral healthy tissues, the digestive, hematopoietic, pulmonary, cognitive and behavioral functions of the organism, as well as of the regenerative potential of tissues [1, 2-11]. However, there are several problems that must be solved before this treatment modality can be translated safely in clinical settings [2]. In this connection, at present there is an increasing interest in studying the biological effects of irradiation with ions and particles using ultra-high dose rates.

Radiotherapy of tumors with hadron beams is being increasingly used owing to their beneficial energy deposition profile (spread-out Bragg peak, SOBP), which spares healthy tissues in the vicinity of the tumor, and to additional positive effects within the tumor associated with high linear energy transfer (LET). Proton radiotherapy, as distinct from irradiation with heavy ions, e.g., carbon ions, makes it possible to effectively preserve healthy tissues in the region after the Bragg peak [12] and even in the plateau region upon FLASH irradiation, as was shown on laboratory mice [8].

The researchers of the FLASH effect believe that further experiments are needed to confirm the presence/absence of the protective effect for healthy tissues and to determine the dose limits and requirements for the practical application of the FLASH irradiation regime, considering that the few experimental studies performed used different types of radiation with different LET and different location in the Bragg peak for hadrons and ions [2]. Since the exposure to protons using different regimes induces multidirectional responses, it is evident that the research should be continued, among other things, with irradiation of healthy tissues.

A biological model used in the present study was fertilized eggs of Japanese quail (*Coturnix japonica*), which were irradiated on the linac at the Institute for Nuclear Research, Russian Academy of Sciences (INR RAS) both in the SOBP and plateau in three modes corresponding to the mean dose rates: SPLASH, D ∼ 10^6^ Gy/s; FLASH, D ∼ 100 Gy/s; and CONV, D < 1 Gy/s. The term SPLASH (single-pulse FLASH) in this work denotes a very high dose rate of about 10^6^ Gy/s. The model has the following advantages: (1) eggs are a model of organogenesis and tissue differentiation; (2) the small sizes of a quail egg make it possible to bring the position of the egg into coincidence with the SOBP and more precisely control the dose; (3) it is possible to assess irradiation-induced changes on the organism level at remote terms (at the stage of embryogenesis and in the postembryonal period); (4) it is possible to synchronize the development of bird embryos due to simple warming of eggs, which substantially facilitates the work and does not require the urgent delivery of classical actively developing biological models of development such as *Danio rerio*; (5) it is possible to facilitate the delivery of the dose in the Bregg peak due to short-term immersion of incubated eggs in the aqueous medium phantom without destroying the object and its normal development; and (6) a high degree of similarity of the embryogenesis of birds to that of man (embryogenesis of higher vertebrates differs from the anamniotic models of embryogenesis: the zebrafish and xenopus).

## 2. Materials and Methods

### 2.1. Animals and cells

Fertilized eggs of the Japanese quail (*C. coturnix japónica*) obtained from the LLC Shepilovskaya Poultry Farm were used [13]. The eggs were stored for no longer than 1 day at a temperature of +15^о^С and then placed in an incubator with programmable temperature, humidity and egg turning automatic control for the proper development of embryos. On the 3^rd^ day, eggs containing living embryos at an identical stage of development were selected using an ovoscope. Irradiation was carried out between the 5^th^ and 6^th^ day of incubation. The incubation procedure was the same for irradiated and unirradiated eggs. The programmable incubator (Spektr-84-01, RF) ensured the maintenance of the preset parameters: the temperature +37.8°С (days 1-7), +37.6°С (days 8-14), +37.4°С (days 15-18); humidity 75% (1 day), 65-75% (days 2-9), 40-45% (days 10-14), 65-75% (days 15-18); ventilation frequency was twice a day for 5 min (days 3-7), twice a day for 15-30 min (beginning on the 8^th^ day); egg turning was carried out once every six hours until the 17^th^ day of incubation; on the last day of incubation, eggs were not turned. A day prior to the hatching of chicks, eggs that received a similar dose in the same mode were placed in one brooder unit. Chicks had free access to water and food throughout the time of the experiment except when behavioral tests were performed. The analysis for each group of irradiated eggs was performed in a similar way. Chicks were killed by decapitation.

Splenocytes were obtained using a 9-week-old outbred SHK mouse male maintained under standard conditions of the vivarium at the Institute of Theoretical and Experimental Biophysics, Russian Academy of Sciences (ITEB RAS, Pushchino, RF) at a temperature of 20 ± 2^о^С, natural daylight, and free access to water and food. The mouse was killed by cervical dislocation. The spleen tissue was squeezed through a nylon mesh, the cell suspension was diluted with phosphate-buffered saline (PBS), and splenocytes were concentrated by centrifugation in a ficoll‒verografin gradient as described [14]. All manipulations with model animals (mice, embryos, chicks) were performed in accordance with the Helsinki Declaration of 1975 (revised in 1983) for the care and use of laboratory animals and the recommendations of the Biological Safety and Bioethics Commission at ITEB RAS (protocols nos. 20/2021 and 23/2023).

### 2.2. Irradiation facility and irradiation technique

Eggs were irradiated on the INR linac, (Troitsk, Moscow), which provides a unique possibility to irradiate biological objects with accelerated protons in a SPLASH mode with mean D of up to 1 MGy/s [15]; it has an energy range of 105-269 MeV, pulse frequencies of 1-100 Hz, pulse duration of 0.3-100 µs, and pulse current from 0.1 µA to 10 mA. Three irradiation modes were used: <u>s</u>ingle-<u>p</u>ulse FLASH mode (abbreviated as SPLASH) with Ḋ ∼10^6^ Gy/s; the standard FLASH mode with Ḋ ∼100 Gy/s; and the CONV mode with Ḋ < 1 Gy/s. Dosimetry was performed using calibrated EBT-XD radiochromic films (GafChromic, USA) and a self-made Cherenkov monitor [16]. In addition, in the CONV mode, doses delivered to eggs were additionally controlled using an PTW Advanced Markus 34045 ionization chamber with an electrometer PTW Multidos. Eggs fixed in a self-made holder were placed in an water phantom and irradiated for short time in different modes. Immediately after the exposure, eggs were taken from the container and placed in an incubator. The location of eggs during the irradiation by a proton beam is shown in Fig. 1.

**Fig. 1.**
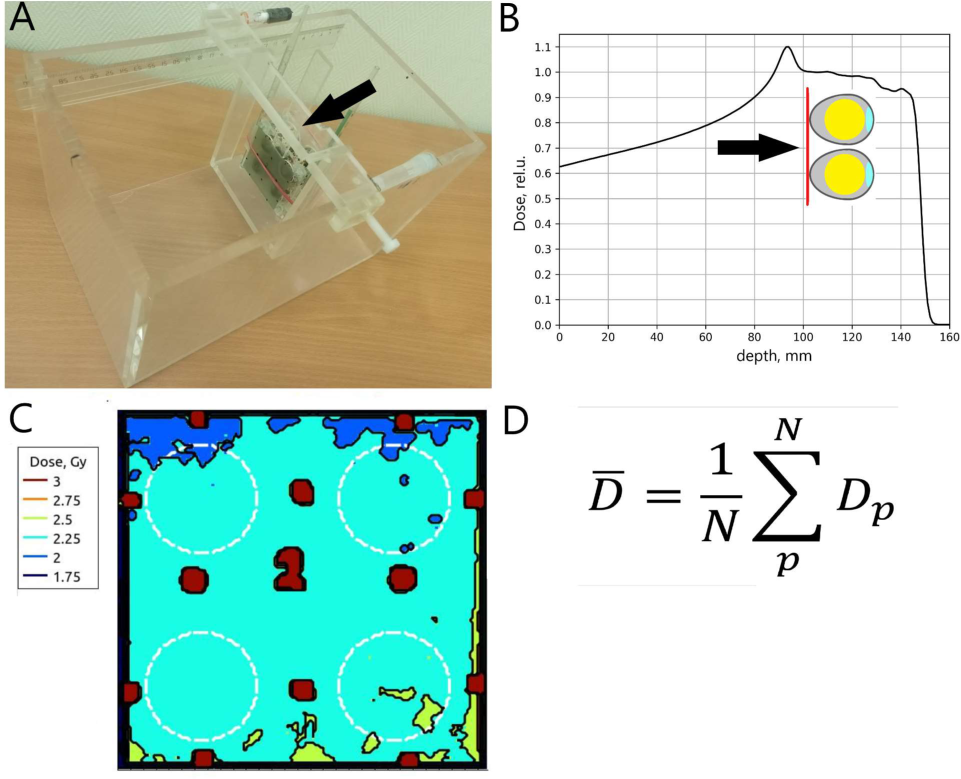
A scheme of the location of eggs for irradiation with protons. (A) A photograph of a water phantom and the location of eggs in the egg holder (shown by the arrow). (B) The location of eggs relative to the SOBP and of the radiochromic film relative to the eggs. The radiochromic film is shown by the arrow. (C) 2D isodose distribution; computer calculations based on the data of the radiochromic film. (D) The calculation of mean doses is based on 2D isodose distribution.

Eggs were divided into 12 groups, with an average of 8-12 eggs in each group. In the control group (unirradiated eggs), 20 eggs out of 40 were opened on the 9^th^ day, and from the remaining 20 eggs, chicks appeared on the 18^th^ day. Control eggs were subjected to the same physical manipulations (transportation, transfer in the incubator) as irradiated. Irradiated eggs were divided into the following groups according to the median dose: three groups were irradiated in the SOBP, the SPLASH mode, with the median doses of 2, 4, and 8.5 Gy; two groups were irradiated in the SOBP, the FLASH mode, with the median doses of 4 and 8.5 Gy; three groups were irradiated in the SOBP, the CONV mode, with the median doses of 4, 6, and 8.5 Gy; two groups were irradiated in the plateau region, in the SPLASH mode, with the median doses of 6 and 14 Gy; and one group was irradiated in the plateau region in the CONV mode with a dose of 7 Gy. After irradiation, the eggs were maintained in the incubator. Chicks hatched from irradiated eggs on the 18^th^ day, with the hatching time of about half a day. In the groups of animals irradiated with 4 and 8.5 Gy, the parameters (the body weight and length, cytogenetic anomalies, and activity of chick) in five (in rare cases four) survived 1-day-old chicks visually having no developmental deviations were analyzed. Each group of irradiated and control chicks was maintained in a separate chamber under similar lighting, humidity, and temperature conditions. An analysis of the body weight and length, cytogenetic anomalies of embryos and chicks, as well as the activity of chicks was performed for the groups of animals irradiated with a similar dose (M ± SD). For embryos, the dose was: 8.4 ± 0.5 Gy in the group irradiated in the SPLASH mode, 8.5 ± 0.4 Gy in the group irradiated in the FLASH mode, 8.8 ± 0.8 Gy in the group irradiated in the CONV mode; for chicks: 3.7 ± 0.5 Gy for animals irradiated in the SPLASH mode, 4.0 ± 0 Gy in the group irradiated in the FLASH mode, and 3.7 ± 0.4 Gy in the CONV mode. The analyzed biological parameters were compared after irradiation mode-base grouping due to no dose dependence for body length and weight was observed for dose interval 7,5-9,5 Gy in case of each irradiation mode in SOBP (data not shown).

Mouse splenocytes were irradiated with protons and X-rays at doses of 5 and 10 Gy after their immobilization in agarose on glass slides followed by the lysis procedure (see the subsection Comet assay). Slides with lysed cells were glued with agarose to the bottom of Petri dishes 35 mm in diameter. During irradiation, the free space of Petri dishes was filled to the top with the lysis solution to prevent the drying of slides. Before irradiation, Petri dishes (three pieces per package) were hermetically sealed in cellophane and fixed with holders in a water phantom. On the outside of the bottom of each dish, there was an EBT-XD radiochromic film (GafChromic, USA). The slides were irradiated with protons in triplicate in the SOBP in the SPLASH, FLASH, and CONV modes. For comparison, part of these slides in the presence of the lysing solution was irradiated on a RUT-250-15-1 X-ray facility of the Center of Collective Use «Radiation sources» at the Institute of Cell Biophysics of the Russian Academy of Sciences (ICB, RAS, Pushchino, Russia) at the following parameters: dose rate 1.12 Gy/min, voltage 200 kV, current 20 mA, filters 1 mm Al and 1 mm Cu, focus distance 37 cm. Dosimetry was carried out by the method of Fricke in the presence of benzoic acid and also using an RFT-Dozimeter VA-J-18 instrument.

### 2.3. Comet assay

The alkaline variant of the comet assay was performed as described in [17] with minor modifications [18]. Briefly: cells were immobilized in low-melting agarose on slides; after the solidification of agarose, they were lysed in a solution containing 2.5 М NаС1, 0.1 М EDTA, 0.01 М Tris-НСl, рН 10, and 1% Triton Х-100 followed by denaturation and electrophoresis at рН > 13, washing, staining of slides, capturing of microscopic images and their analysis. The DNA damage was estimated using two parameters: the percentage of DNA in the comet tail (%TDNA) and the comet tail length (TL) in pixels. For each experimental point, three slides were used, with no less than 100 cells being analyzed on each.

### 2.4. Preparation and staining of blood smears

Blood from chicks was collected during decapitation, and embryonic blood was taken from the yolk sac vascular network after the removal of the eggshell. Blood smears prepared by the standard method on slides [19] were air-dried, fixed in 96% ethyl alcohol for 20 min, stained with azurin eosin by the method of Gimsa-Romanovsky, washed with PBS, рН 7.2, dried, and analyzed. Then, the smears were examined under a bright-field light microscope (LOMO, RF) at a magnification of ×900. Photographs were taken on a Leica DM2000 light microscope with a Leica DFC425 color camera at a magnification of ×1000. In smears, the number of erythrocytes with three types of anomalies were quantified: erythrocytes without nuclei, with micronuclei, and with two nuclei (a figure-of-eight nucleus). The total number of erythrocytes in the field of vision was counted. The total number of analyzed cells was no less than 3000 per smear. The percentage of cells with anomalies was calculated.

### 2.5. Analysis of chick mobility

The activity of chicks was assessed using the open field method with modifications that correspond to the common behavioral features of chicks of brood birds [23, 24]. Groups of four to five chicks 24 h after hatching from irradiated eggs were taken in the experiment. Chicks were placed in a square arena (40 cm long, 55 cm wide, wall height 30 cm) covered with graph paper. The movements were recorded under natural light using a digital photo/video camera (Olimpus, Japan) secured above the open field arena. Simultaneous staying in the open air of a group of one-day-old chicks reduced the stress and assured a more natural behavior of chicks of brood birds prone to group together [23] The defecation products of chicks were removed between recording series. Video records of movements of similar duration were analyzed beginning from the first frame. The time intervals of video recording were so chosen that chicks had no time to form a group, and the level of stress they developed was not sufficient to send non-stop distress signals. The minimum observation time was 5 min. For each individual, a frequency graph of movement velocities with a bin width of 5 mm/s was compiled. Then, for each velocity interval, the mean and mean-square deviations of the frequency of movement repetitions in a preset velocity range were calculated. To determine the least mobile group, the individual mean-arithmetic velocities of movement in a given time interval were calculated, after which a one-factor dispersion analysis ANOVA was performed for the hypotheses of pairwise statistical correspondence of the mean velocity to the irradiation mode.

The movements of individuals were tracked using the Discriminative Correlation Filter with Channel and Spatial Reliability Tracker (CSRT) from the open computer vision library openCV (https://opencv.org). The CSRT algorithm uses spatial reliability maps to tune the filter to a part of the selected area from the tracking frame, which makes it possible to increase the search region and track nonrectangular objects. When examining a record, it is possible to detect the superposition of objects and continue maintaining the history of movements starting from the moment the tracker made a mistake. A result of the tracker operation at an arbitrary time moment is the coordinates and the sizes of a rectangle that corresponds to the position of the object in the frame in pixel units. Given the number of a current frame F, the coordinates of the rectangle center (x, y) that correspond to the position of a chick in the space, the spatial resolution of image R (pixel/mm), and the frame rate, it is possible to estimate the movement velocity. In this case, the distance traveled by a chick is calculated by the Pythagorean theorem based on the coordinates of two points in a specified range of frame change. One of a CSRT features is the possibility to predict the movement based on the history of movements and estimate the movement velocity. However, when analyzing a change in the coordinates after one frame, a discreteness of the velocity spectrum is observed. To eliminate this effect, the velocity was analyzed in the Δt interval of five frames, which at 60 fps corresponds to one twelfth of a second. For the analysis, several timelines with a step of τ ɛ [0 Δt) were used. The activity of chicks was compared by velocity using the method of Tiemann [24] with some modifications.

### 2.6. Statistical processing

The statistical significance of differences was analyzed using the Student’s *t*-test and the disperse analysis ANOVA. The significance of differences was taken at *р* < 0.05.

## 3. Results

### 3.1. The level of DNA damage in the nucleoids of mice splenocytes exposed to protons is higher than that upon X-irradiation

In preliminary experiments, the level of DNA damage in irradiated mouse splenocytes was determined by the comet assay. In these experiments, only irradiation-induced DNA damage was estimated, without DNA repair. Figure 2 shows the levels of DNA damage according to the criteria of tail length (TL) of comets (Fig. 2A) and %TDNA (Fig. 2B) after *in vitro* exposure to X-rays or accelerated protons in SOBP.

**Fig. 2.**
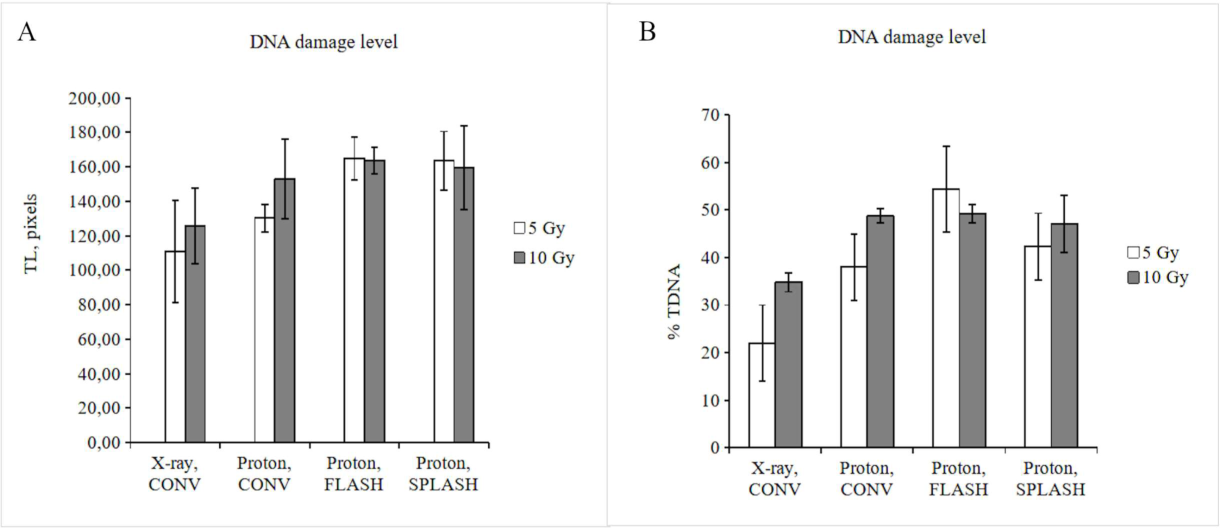
Irradiation with protons in the SOBP induces a higher level of DNA damage in mouse splenocyte nucleoids than X-irradiation *in vitro.* DNA damage was evaluated by the Comet assay using the following parameters: (A) the average comet tail length TL, (B) the average percentage of DNA in the cornet tail ¾TDNA. Groups (bars indicate the mean, M) were formed according to the dose delivery mode and irradiation source (CONV, FLASH, SPLASH, protons; X-rays). The mean was calculated by the values for individual irradiated rnicroglasses; the error bars show the standard deviation SD (M ± SD).

As seen from Fig. 2, the level of damage to nucleoids by protons is higher than after X-irradiation. In the FLASH/SPLASH modes, as opposed to the X-ray irradiation, no dependence of DNA damage on the dose was observed.

### 3.2. Irradiation in the FLASH and SPLASH modes at a dose of 8.5 Gy causes a significant reduction in the damage to embryos

Based on the curves shown in Fig. 2 and the reported data [22-24], the mean doses of 4 and 8.5 Gy were chosen for the irradiation main part of of eggs. Eggs irradiated with doses of 8-14 Gy were opened on the 9^th^ day of development (the number of survived embryos is given in a Table 1), and eggs irradiated with doses of 2-7 Gy were incubated until chicks hatched (Table 2). Unfertilized eggs that were opened on the 9^th^ and 18^th^ days, as well as eggs that ceased to develop on the 1^st^-3^rd^ days after placing into the incubator were excluded from consideration.

**Table 1.** Survival rate of 9-day-old embryos from eggs irradiated with protons in different modes in the dose range of 8-14 Gy.

| Irradiation mode | Irradiation in SOBP | Dose, Gy | Number of irradiated eggs | Number of survived embryos | Note |
| --- | --- | --- | --- | --- | --- |
| SPLASH | + | 8 | 16 | 14 | 2 died early in development |
| FLASH | + | 8 | 15 | 15 | All embryos have poorly developed blood vessels |
| CONV | + | 8 | 14 | 14 |  |
| SPLASH | - | 14 | 7 | 0 |  |

As seen from Table 1, with each mode of irradiation with 8.5 Gy in SOBP, most embryos remained alive after the 9^th^ day; all embryos irradiated with the FLASH mode, showed a weaker filling and development of blood vessels compared with control. After irradiation with 14 Gy in the plateau region, 100% of embryos were dead on the 9^th^ day.

Figure 3 shows the body length (Fig. 3A), and the body weight (Fig. 3B) of 9-day-old embryos from eggs irradiated with 8.5 Gy, as well as the typical images of micronuclei in erythrocytes (Figs. 3C, 3D) and the percentage of erythrocytes with micronuclei (Fig. 3E) detected in embryonic blood.

**Fig. 3.**
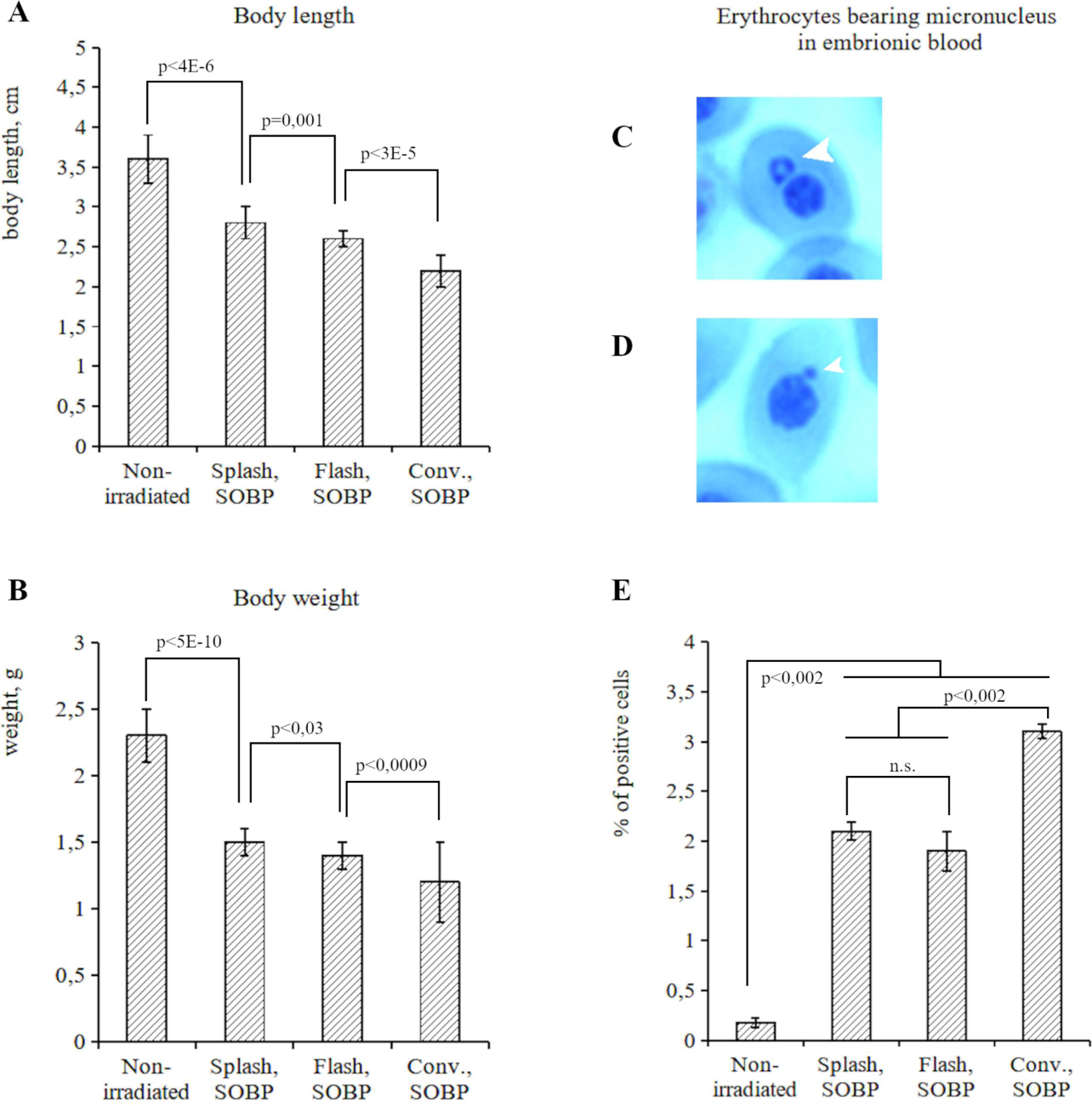
According to the weight, sizes, and the number of erythrocytes with rnicronuclei in 9-day-old embryos from eggs irradiated with a dose of ∼8.5 Gy in the SOBP, the FLASH and SPLASH irradiation modes were found to have the least damaging effect. (A) Body length of the embryo, cm. (B) Body weight of the embryo, g. (C, D) Images of erythrocytes with rnicronuclei; rnicronuclei on the photo are shown with arrow heads. (E) A quantitative analysis of the number of erythrocytes with rnicronuclei in samples. Groups (bars indicate the mean M) were formed according to the dose delivery mode (CONY, FLASH, SPLASH). The mean was calculated by the values for individual embryos; the error bars show SD (M ± SD).

It is seen that, with all irradiation modes, the length and the weight of 9-day-old embryos are smaller than those of nonirradiated. With the SPLASH mode, the maximum body weight and length were observed compared with the other irradiation modes. Significant differences in the average body weight values in groups were observed between SPLASH and FLASH modes (*p* < 0.03) and between the SPLASH or FLASH and CONV modes (*p* < 0.00003 and *p* < 0.0009, respectively). Significant differences in the average body length values were observed between SPLASH and FLASH modes (*p* = 0.001) and between the SPLASH or FLASH and CONV modes (*p* < 0.000002 and *p* < 0.00003, respectively). It was found that, with all irradiation modes, the number of erythrocytes with micronuclei in embryos increased and significantly differed from the control (*p* < 0.05). The maximum number of these erythrocytes was observed after irradiation in the CONV mode, and it was significantly different from that observed in the SPLASH or FLASH modes (*p* = 0.002 and 0.0005, respectively). All chicks that hatched on their own from eggs irradiated in different modes did not differ in appearance from one another and from the control ones.

### 3.3. Cytogenetic anomalies of erythrocytes and the body weight of 1-day-old chicks demonstrate no statistically significant difference between the irradiation modes at a dose of 4 Gy

In Table 2, the data on the survival of chicks hatched on the 18^th^ day from eggs irradiated with protons are given.

**Table 2.**
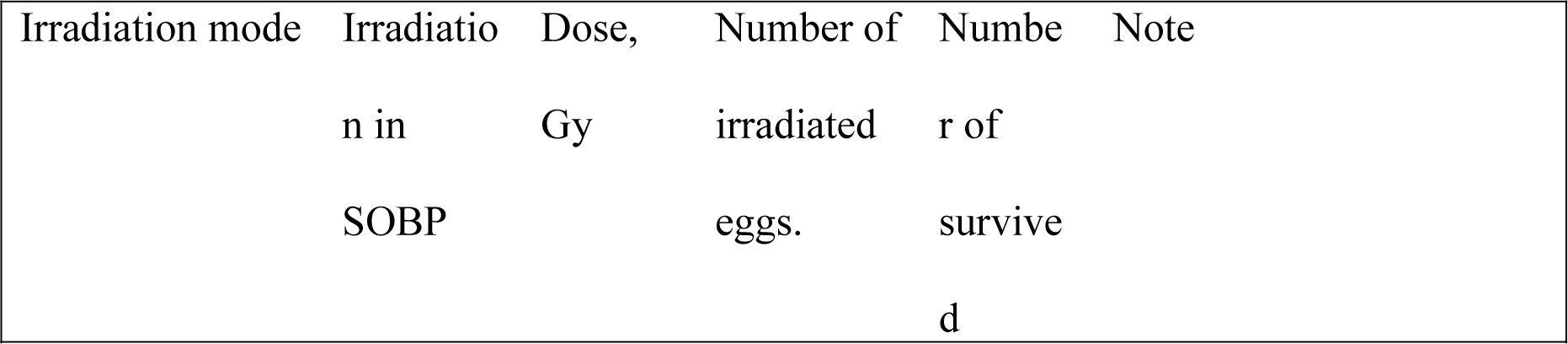
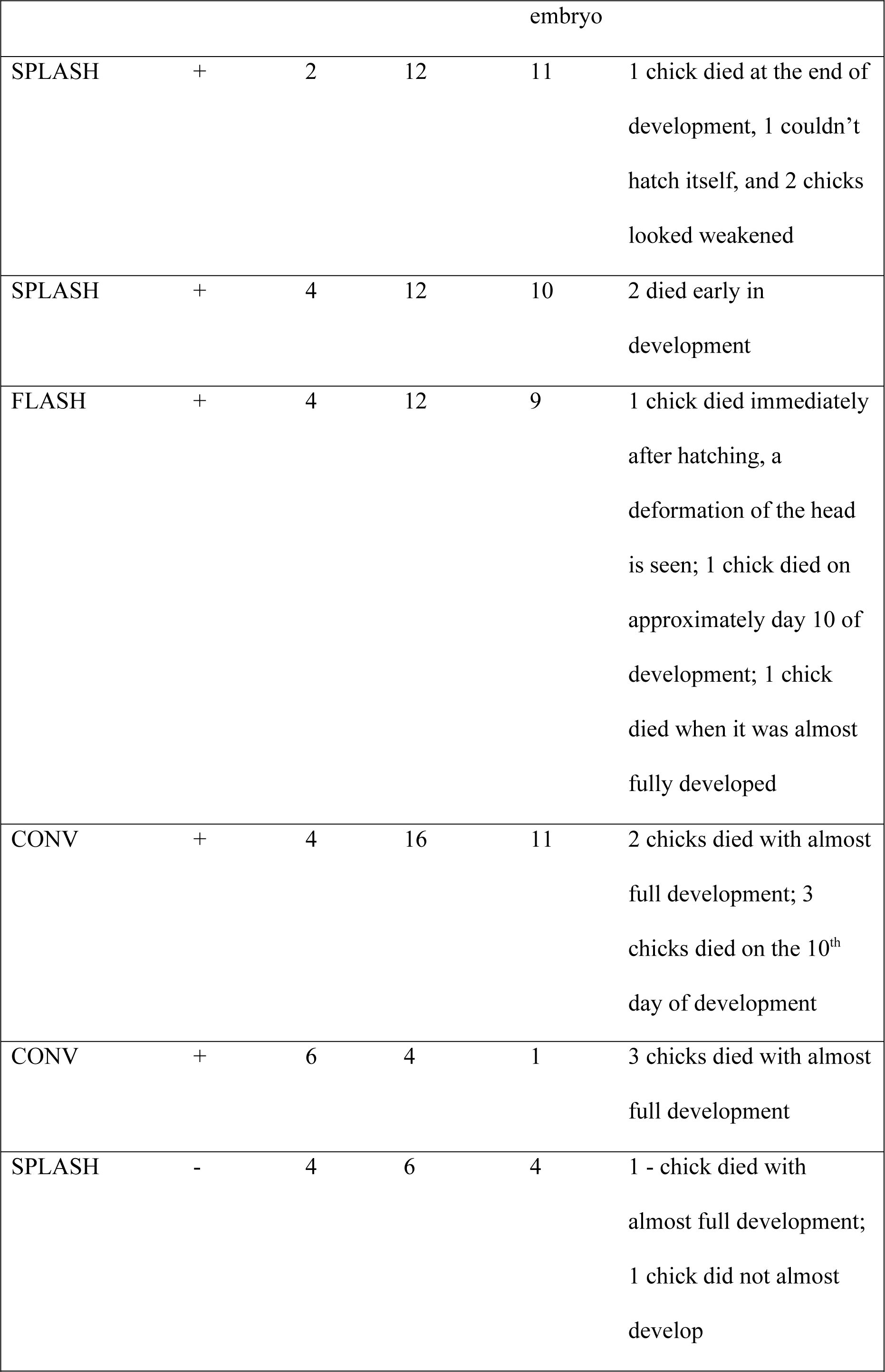
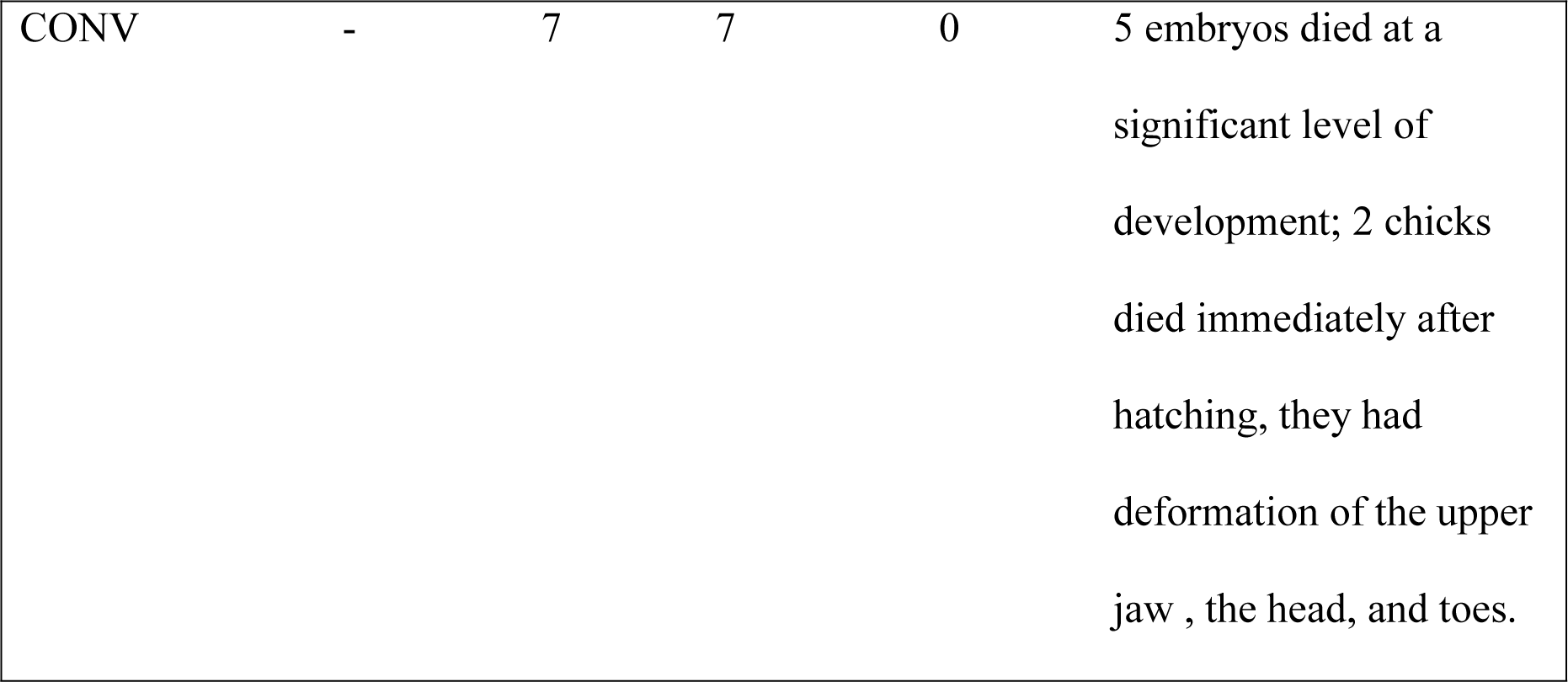
Survival rate of chicks from eggs irradiated with protons in different modes in the dose range of 2-7 Gy.

It is seen that, after irradiation of eggs with 2 Gy in the SPLASH mode (SOBP), about 8% of animals died, and after irradiation with 4 Gy, about 16-17%. After irradiation with 4 Gy in the FLASH and CONV modes, about 25% and 31% deaths, respectively, were recorded. The maximum deaths were recorded after irradiation in the plateau region.

Figure 4 gives the weights of chicks hatched from eggs irradiated with 4 Gy. It is seen that the weight of all survived chicks corresponds to that of control animals and does not statistically differ for different irradiation modes.

**Fig. 4.**
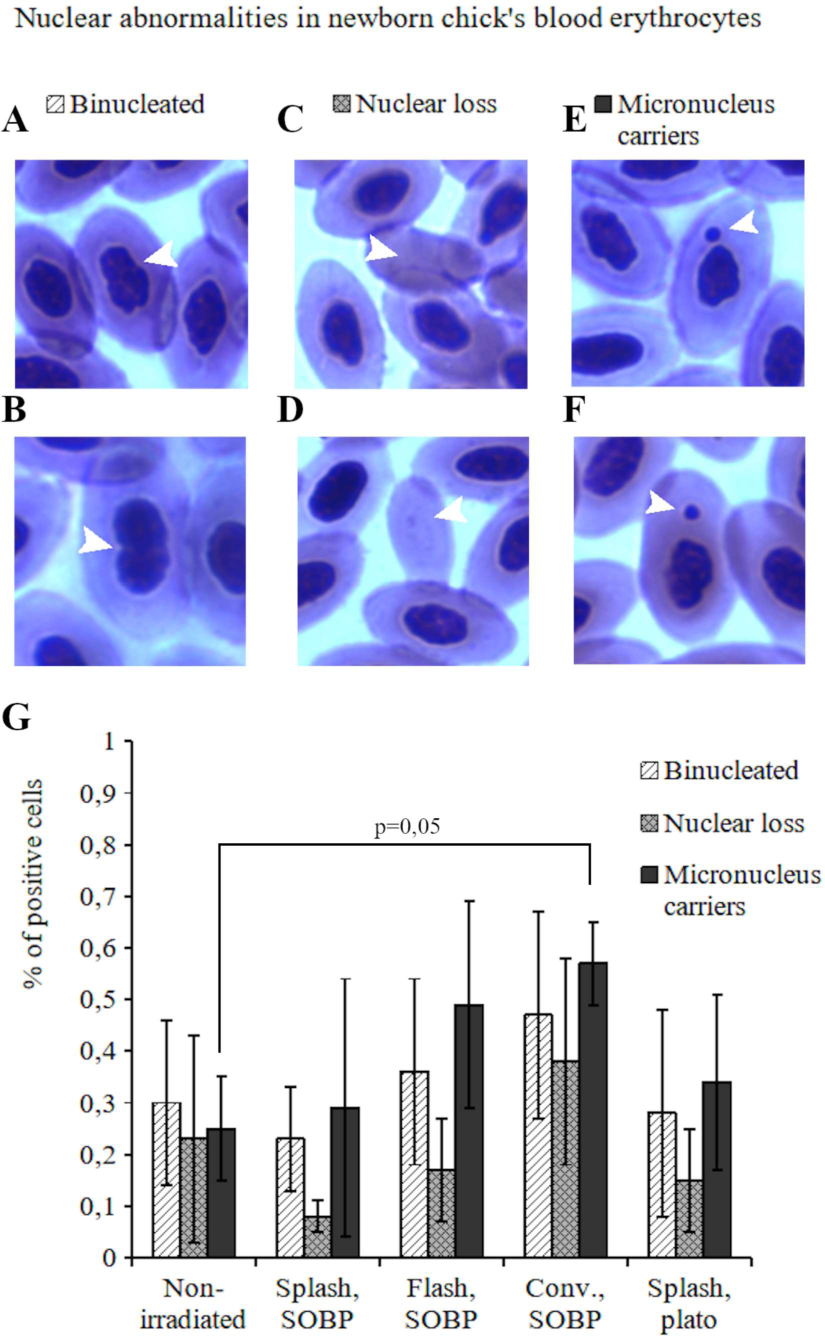
The body weight of survived 1-day-old chicks corresponds to that in the control and does not statistically differ among different irradiation modes. Eggs were irradiated with a dose of ∼4 Gy in the SOBP and in front of it. Groups (bars indicate the mean, M) were formed according to the dose delivery mode (CONY, FLASH, SPLASH). The mean was calculated by the values for individual chicks; the error bars show SD (M ± SD).

Figure 5 shows the counts of anomalous forms of erythrocytes in blood smears from 1-day-old chicks.

**Fig. 5.**
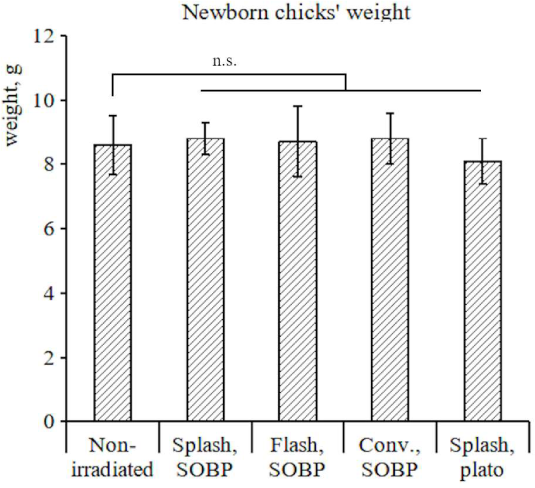
SPLASH and FLASH modes of proton irradiation were found to have the least damaging effect on blood erythrocytes. (A, B) Photos of binucleate erythrocytes, (C, D) photos of anucleate erythrocytes, and (E, F) photos of erythrocytes with micronuclei in 1-day-old chicks irradiated with a dose of ∼4 Gy in the SOBP and in the plateau. Cytogenetical abnormalities are labeled with arrow heads. (G) A quantitative analysis of the number of erythrocytes with cytogenetic anomalies in samples. Groups (bars indicate the mean, M) were formed according to the dose delivery mode (CONV, FLASH, SPLASH). The mean was calculated by the values for individual animals; the error bars show SD (M ± SD).

It is seen that there is no significant difference in the number of binucleate erythrocytes, as well as anucleate erythrocytes and erythrocytes with micronuclei both between the groups irradiated in the SPLASH mode in the SOBP and the plateau as well as in the FLASH mode in the SOBP; also, there is no difference between these irradiation modes and the control. However, an almost significant (*p* = 0.051) increase compared with the control was observed in the number of erythrocytes with micronuclei after the irradiation in the CONV mode (Fig. 5 G).

### 3.4. Chicks from eggs irradiated in the CONV mode are less active than chicks from all other eggs

The results of the analysis of the mobility of chicks are shown in the histogram in Fig. 6. As evidenced by the video, a distinguishing feature of movements of nonirradiated quails is their coming together to form groups followed by coming apart. A minimum time until the grouping was 33 s (1978 frames).

**Fig. 6.**
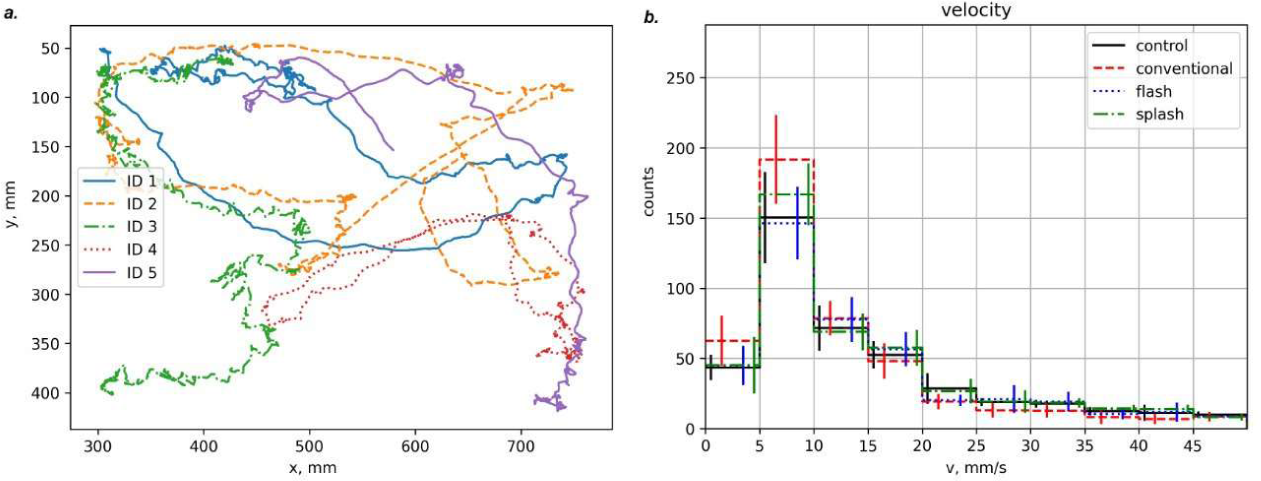
Chicks from eggs irradiated in the CONV mode are less active than chicks from nonirradiated eggs and eggs irradiated in the SPLASH and FLASH modes. (A) An example of the recognition of individual tracks of five chicks located simultaneously in the open field. (B) A histogram of the mean velocity of the movement of 1-day-old chicks hatched from the eggs irradiated with a dose of ∼4 Gy. Bars indicate the mean (M) for the ranges of movement speed (the abscissa); on the ordinate is the frequency of occurrence of a particular velocity within the velocity range. Groups (bars indicate the mean, M) were formed according to the dose delivery mode (CONV, FLASH, SPLASH). The mean was calculated by the values for individual chicks; the error bars show SD (M ± SD).

As seen from Fig. 6, the rate of movements of chicks hatched from eggs irradiated in the SPLASH and FLASH modes is comparable with that of control birds. Chicks irradiated in the CONV mode are less mobile than chicks from control eggs (*p* = 0.002) and eggs irradiated in the FLASH (*p* = 0.006) and SPLASH modes (*p* = 0.019).

## 4. Discussion

In this work, we estimated some characteristics of the state of embryos on the 9^th^ day (middle of the development period) and of chicks 24 h after hatching from incubated eggs irradiated with protons at different dose rates (Figs. 3-6). To choose optimal doses, the level of radiation-induced DNA damage (%TDNA, TL) in the nucleoids of mouse splenocytes was preliminarily estimated (Fig. 2). Splenocytes were taken as an object that is commonly used in radiobiology to evaluate the general level of DNA damage in radiation-sensitive cells. One mouse was taken for obtaining the splenocytes to minimize the data scattering due to individual features of the animals in a group. In irradiated nucleoids, no DNA repair occurs; therefore, it is possible to estimate the level of damage induced only by ionizing radiation. As was to be expected, %TDNA and TL after exposure to protons were greater than those after X-ray irradiation (Fig. 2). A comparison of the effects of proton beams (60 MeV) and X-rays (250 kV) in the dose range of 0.3–4.0 Gy on human peripheral blood lymphocytes showed that the probability of cell death after irradiation with protons is higher than after X-rays exposure [24]. The comparison did not reveal the dependence of the DNA damage on dose in the FLASH/SPLASH modes (Fig. 2) in the dose range of 5-10 Gy due to the saturation effect and a decrease in the sensitivity of the method under the experimental conditions used [17]; therefore, a dose of ∼8.5 Gy was chosen as a larger dose for examining the eggs on the 9^th^ day (it was expected that, with a dose of ∼8.5 Gy, most embryos would be incapable of hatching on the 18^th^ day due to the severity of damage) and a dose of ∼4 Gy to discover significant differences by the end of incubation (on the 18^th^ day, chicks were expected to hatch according to their normal development). It was assumed that the DNA damage in these chicks would be sufficient to be detected and to make conclusions on the differentiated impact of different dose rates.

Most embryos from eggs irradiated with a dose of 8.5 Gy were alive on the 9^th^ day (Table 1); the length and weight of their bodies in all radiation modes were significantly less than those of nonirradiated (Figs. 3A, 3B). It is evident that the reduction in these parameters could hamper hatching in case the embryos remained alive, since the weight of an embryo and, in the final analysis, of a chick is an important parameter for egg hatching; during the formation of the skeleton, calcium transfers from the shell into the embryo so that the shell thins and the hatching is facilitated. According to the criteria of body weight and length as well as the number of erythrocytes with micronuclei, the SPLASH/FLASH modes were the least harmful: these parameters statistically significantly differed from those recorded after irradiation in the CONV mode (Fig. 3). It should be noted that the percentage of erythrocytes with micronuclei in embryos was higher than in chicks (Fig. 3E and Fig. 5G), which may be due to both the completion of the development of a chick and the use of different doses, 4 Gy for chicks and 8.5 Gy for embryos.

In hatched chicks, no marked differences in appearance from intact animals were found with all irradiation modes, which agrees with the data obtained after the X-ray exposure of hen eggs. Previously no drastic external abnormalities among chickens hatched from hen eggs X-irradiated at a dose of 5 Gy was found; however, late hatching with growth retardation and a lower body weight compared with chicken hatched from irradiated (0.5 Gy) and unirradiated eggs were noted [22]. On the average, a later hatching of chicks from eggs irradiated in the CONV mode, but not in the FLASH and SPLASH modes, was also observed in the present work compared with the control. It is seen that significant differences in the weight and length of embryos were dependent on irradiation mode (Figs. 3A, 3B), whereas the weight of chicks was not (Fig. 4). Presumably, the differences from irradiated hen eggs are due not only to a lower dose (4 Gy in our study compared with 5 Gy) but also by the other type of exposure as well as some features in the development of quail embryos. It may be suggested that the differences in the weight and length, observed in embryos on the 9^th^ day under different irradiation modes, are not only due to a high radiation dose (8 Gy) but partially also to the regulation of development rate caused by different irradiation modes. Probably, the retardation in the development is related to a lower activation of signaling through wider activating the expression of the ligand of the transforming factor beta (TGF-β 1), which is characteristic of the CONV irradiation mode in contrast to FLASH and the nonirradiated control. A large imbalance of the TGF-β-mediated signaling after the CONV irradiation can also cause a retardation in tissue differentiation and the weight gain of the embryo [1, 25-27]. Despite the lower dose of 4 Gy, some embryos died on the 18^th^ day under all irradiation modes (Table 2). A lower death incidence was observed with the SPLASH mode (approximately 16-17%); after irradiation in the FLASH and CONV modes, 25% and 31% embryos died, respectively. The greatest number of deaths was recorded after irradiation outside the SOBP, which can be explained by a higher flow density of particles and significantly more abundant point mutations in DNA since the integrity of the whole genome is important for embryogenesis and cell differentiation. In turn, most fully differentiated somatic cells in adults may show a better resistance to these point mutations. A high rate of cell proliferation in embryonic tissues may partly compensate for the loss of cells with more complex and unrepairable DNA lesions caused by protons in the SOBP. At the higher dose of 7 Gy in the CONV mode (the plateau region), in two chicks that hatched from irradiated eggs and died immediately after hatching, a deformation of the head, underdevelopment of toes and the upper jaw were observed; however, since the events were few in number and the sampling size was small, no conclusions about the quantitative significance of the results could be made. An analysis of defects in the formation of limbs after the X-irradiation of their buds in hen embryos showed that, 3.5 and 5.5 days after the start of incubation of eggs, predominantly the proximal and distal wing buds, respectively, are damaged [28]. The authors associate this with the death of chondrocyte precursors, a definite amount of which is necessary for the condensation of skeletal elements [29]. At earlier stages, the condensation of more proximal limb parts occurs, and at later stages the condensation of more distal limb parts takes place, which just explains different sensitivity to irradiation. After the condensation of limb buds, the radioresistance of their constituent cells increases, which may ensure a better preservation of the elements of limb buds. On the other hand, during the condensation of buds, an imbalance of the differentiation factors (e.g., TGB-β signaling) may occur. The defects in the development of hind limb toes, detected in embryos irradiated in the CONV mode in present work, are similar to the damage to distal limb parts, observed in chicken embryos X-irradiated in the same mode after 5.5 days of development [28]. The data obtained in the present work partially favor the application of the FLASH regime in the therapy of bone and cartilage cancers as a gentler mode; however, its effectiveness against cancer cells has yet to be established [30].

It is known that there are more than 20 abnormalities of nuclei in the erythrocytes of birds [31]. The minimum numbers of all three types of erythrocyte abnormalities revealed in this work were recorded using the SPLASH mode both in the SOBP and the plateau region, as well as the FLASH mode (Fig. 5G). Since the data for these abnormalities had large standard deviations, it was impossible to establish any statistically significant difference between the measured percentage and the control in each group as well as among the groups. It can only be said that there is a tendency for an increase in the number of binucleate and anucleate erythrocytes, and erythrocytes with micronuclei in chicks from eggs irradiated in the SOBP in the CONV mode compared with the SPLASH/FLASH modes and the control. The modest effects on the accumulation of cytogenetic abnormalities are probably associated with the radioresistant function of the antioxidants of the yolk sac, which occupies a considerable volume of the egg on the 5^th^/6^th^ day. The large standard deviations of the data in the groups are probably due to great variations in the numbers of erythrocytes with abnormalities in the norm: the number of erythrocytes with micronuclei (staining with azure–eosin by Romanovsky–Giemsa) in Japanese quails was found to vary from 0.82 to 2.83% in females and from 0.58 to 1.83% in males [32]. The heterogeneity of a macrobeam at the level of its microorganization/density fluctuations may also play a role. If this is the case, the dose received by individual hemopoietic cells would differently damage these cells and thus differently affect the intermediate stages of the development of the chick embryo (the 9^th^ day of development). An indirect support for this is the bystander effect detected in experiments with selective irradiation of the nucleus and the cytoplasm and during the random distribution of proton microbeams in the nucleus and the cytoplasm of MRC-5 human lung fibroblasts. In these experiments, the increase in the yield of micronuclei and ROS was observed not only in irradiated, but also in neighboring nonirradiated cells, the differences in the formation of micronuclei being dependent on whether the nucleus or the cytoplasm was damaged [33]. Mitochondria, being the main regulators of apoptosis and inflammation, may play an important role in the FLASH effect, in which, the degree of apoptotic and inflammatory responses significantly decreases compared with the CONV mode [34, 35]. It is likely that similar processes also occurred in our experiments during the development of eggs irradiated at a high dose rate.

In this work, the average frequency of movements of chicks in definite speed ranges, as an indicator of the functioning of the organism was also analyzed (Fig. 6). When moving in an open field of specified sizes, chicks could see each other, so that there was no need to send distress signals typical of brood chicks. Chicks from eggs irradiated in the SPLASH/FLASH modes moved at an average speed comparable with that of control animals. Chicks from eggs irradiated in the CONV mode were less mobile (the mean speed 5-10 mm/s) and showed a greater tendency to gather in groups than chicks from intact eggs and eggs irradiated in the other modes. Since the temperature in the room is lower than in the brooder, it is clear that chicks can gather in groups to get warm, which may indirectly indicate the disturbance of energy metabolism, mitochondrial dysfunction [35], and cognitive dysfunction, which show up in a lower inclination for search behavior. Previously, a greater preservation of nervous tissue during irradiation in the FLASH modes has been shown in irradiated mice and rats [4, 9, 10]. The differences identified in this study indicate that the CONV mode has a stronger damaging effect, which probably affects both energy metabolism and cognitive functions.

Thus, the irradiation of eggs with proton beams in the Bragg peak in the high-dose-rate mode is less damaging to healthy tissues and the embryogenesis of amniotes (quail), which so far has been shown only for the anamniote *Danio rerio* fish [4, 36].

## 5. Conclusions

The fetus exposed to irradiation is not a typical model for assessing the radiotoxicity of accelerated protons. Nevertheless, the examination of embryogenesis after radiation exposure allows a better understanding of the toxic effect of protons on genome integrity in a developing organism compared to a mature one. This increases the number of targets for radiation and adds to the value of this biological model. Previously, in a model of the fish *D. rerio*, the response to irradiation with accelerated electrons and accelerated protons in the plateau region in the FLASH mode was studied [4, 13]. It was found that, at high doses in the FLASH mode, a reduction in morphological alterations after exposure to electrons (10-20 Gy) and a decrease in pericardial edema after irradiation with protons (22-25 Gy) occur. The data on the radiation-induced impairment of embryogenesis of Japanese quail obtained in this work are fundamentally new: 1) an amniotic development model was used; 2) most of the effects have been studied in the Bragg peak, which is important specifically for hadron radiotherapy; and 3) the effects were observed both during embryonic development and immediately after its completion. Based on a number of criteria (the weight of irradiated embryos and day-old chicks, body length, the level of cytogenetic abnormalities of red blood cells and the behavior of one-day-old chicks), it was found that exposure to protons in the Bragg peak in the FLASH and SPLASH modes exerts significantly lower radiotoxic effects on healthy tissues compared with the CONV mode.

### Funding

This study was supported by the Russian Scientific Foundation (project no. 22-25-00211).

## Author Contributions

**Conceptualization**: Elena A. Kuznetsova, Ivan A. Yakovlev, Segrey I. Glukhov, Sergey V. Akulinichev

**Data curation:** Elena A. Kuznetsova, Ivan A. Yakovlev, Segrey I. Glukhov, Sergey V. Akulinichev

**Formal analysis:** Elena A. Kuznetsova, Ivan A. Yakovlev

**Funding acquisition:** Elena A. Kuznetsova, Segrey I. Glukhov, Sergey V. Akulinichev

**Irradiation:** Sergey V. Akulinichev, Yury K. Gavrilov, Dmitry A. Kokontsev, Ivan A. Yakovlev, Valeriia V. Martynova

**Animal work:** Elena A. Kuznetsova, Segrey I. Glukhov, Valeriia V. Martynova

**Behaviour analysis:** Segrey I. Glukhov, Ivan A. Yakovlev

**Methodology:** Elena A. Kuznetsova, Ivan A. Yakovlev, Segrey I. Glukhov

**Project administration:** Sergey V. Akulinichev, Segrey I. Glukhov

**Software:** Elena A. Kuznetsova, Ivan A. Yakovlev

**Supervision:** Sergey V. Akulinichev, Segrey I. Glukhov

**Visualization:** Elena A. Kuznetsova, Ivan A. Yakovlev, Segrey I. Glukhov, Sergey V. Akulinichev

**Writing – original draft:** Elena A. Kuznetsova, Segrey I. Glukhov, Ivan A. Yakovlev

**Writing – review & editing:** Elena A. Kuznetsova, Ivan A. Yakovlev, Segrey I. Glukhov, Sergey V. Akulinichev

## Abbreviations

CONV: conventional dose rate (< 1 Gy/s)
FLASH: ultra-high dose rate (∽100 Gy/s)
SPLASH: single-pulse flash dose rate (∽10^6^ Gy/s)

## Acknowledgments

The authors are grateful to N. P. Sirota for providing mini glasses and agarose mini gel mounting facilities and to G. V. Merzlikin for help with irradiation of eggs.

